# Cryo-EM evidence for a common factor in Alzheimer’s and other neurodegenerations

**DOI:** 10.1101/2025.05.13.653829

**Authors:** Leslie R. Bridges

## Abstract

In the last seven years, cryo-EM maps of neuropathological fibrils from Alzheimer’s disease and other neurodegenerations have been released by various authors^1–44^. The first publication^11^ noted an unknown component coordinating with lysine residues in the protein, a finding recapitulated in many succeeding studies. Previous authors have emphasized difficulties in analysing this component^12,20,28,33,43,45^, but current findings, using powerful visualisation software UCSF ChimeraX^46^ on all publicly available maps^1–44^, indicate that the issue is tractable. Lysine-coordinating extra densities have common features, including a Y-shaped substructure, suggestive of a molecular factor in common, in neuropathological fibrils from a wide range of neurodegenerations and involving misfolded proteins beta-amyloid^10,35^, alpha-synuclein^27,37,39,41^, prion protein^17^, tau^1,5,7,8,11,12,15,16,19,22–26,29–33,35,43^ and transmembrane protein 106B^5,9,18,20,24,28,36,44^. A similar component, albeit in non-lysine environments, was found in neuropathological fibrils involving TAR DNA-binding protein 43^2,3^ and TATA-binding protein-associated factor 15^36^. The results suggest the existence of a common molecular factor, a predominantly anionic polymer, linking these diseases and raising the possibility of a unitary basis for Alzheimer’s and other neurodegenerations. Based on evidence here, RNA is a feasible candidate for this putative common factor. Such findings raise the possibility of new diagnostic tests and treatments for these devastating diseases in the future.

---

Alzheimer’s disease (AD), Parkinson’s disease (PD), amyotrophic lateral sclerosis (ALS) and other neurodegenerations share many overlapping clinical and pathological features^47,48^. Until now, however, there has been no evidence of a common underlying molecule or fundamental mechanism. Were such to exist, it would likely revolutionise approaches to diagnosis and treatment. Advances in cryo-EM have made it possible to examine neuropathological fibrils from the brains of patients with AD, PD, ALS and other neurodegenerations at near-atomic resolution. The first of these studies in 2017^11^ made the surprising observation of a component other than protein, opposite lysines 317 and 321, in tau fibrils from a patient with AD. A total of 161 cryo-EM maps of neuropathological fibrils from brains of patients with a wide variety of neurodegenerations have now been published^1–44^ and many have lysine-coordinating extra densities (LCEDs) containing an unknown component. The obvious question that arises is whether the unknown component is the same in all LCEDs, acting as a missing link between these diseases. The authors of previous studies have stopped short of answering this question, on the basis that LCEDs lack sufficient detail^12,20,28,33,43,45^.

The aim of the present study was to examine all cryo-EM maps of human neuropathological fibrils ex vivo from the Electron Microscopy Data Bank (EMDB)^49^, alongside matching atomic models from the Protein Data Bank (PDB)^50^, using powerful visualisation software, UCSF ChimeraX^46^, and techniques of nano-dissection, contour mapping and marker placement. LCEDs were evaluated for strength, morphology and chemical environment, to see whether they are consistent with a molecular factor in common.

## The nature of LCEDs

A total of 161 cryo-EM maps were examined^1–44^, involving 7 proteins, and over 30 neurodegenerative diseases (see Methods and Extended Data Table 1). Eighty-seven of 141 assessable maps (62%) had LCEDs. A total of 233 LCEDs were found, in diverse neurodegenerative diseases, including AD^1,7,11,12,19,22,25,31,32,35^, argyrophilic grain disease (AGD)^33^, amyotrophic lateral sclerosis/parkinsonism-dementia complex (ALS/PDC)^24^, chronic traumatic encephalopathy (CTE)^8^, corticobasal degeneration^1,43^ (CBD), Down’s syndrome^10,15^, dementia with Lewy bodies (DLB)^5,39^, frontotemporal dementia and parkinsonism linked to chromosome 17 (FTDP-17)^30^, atypical frontotemporal lobar degeneration (aFTLD)^36^, frontotemporal lobar degeneration-TDP (FTLD-TDP)^5,20^, Gerstmann-Sträussler-Scheinker disease (GSS)^16,17^, globular glial tauopathy (GGT)^33^, juvenile-onset synucleinopathy (JOS)^41^, limbic-predominant neuronal inclusion body 4R tauopathy (LNT)^33^, multiple system atrophy (MSA)^27,28,37^, multiple system tauopathy with presenile dementia (MSTD)^18^, normal elder^9,44^, Parkinson’s disease (PD)^39^, primary age-related tauopathy (PART)^32^, Pick’s disease (PiD)^29^, progressive supranuclear palsy (PSP)^5,33^, prion protein-cerebral amyloid angiopathy (PrP-CAA)^16^, subacute sclerosing panencephalitis (SSPE)^23^ and vacuolar tauopathy (VT)^26^, variously involving the misfolded proteins beta-amyloid (Aβ)^10,35^, alpha-synuclein (α-syn)^27,37,39,41^, prion protein (PrP)^17^, tau^1,5,7,8,11,12,15,16,19,22–26,29–33,35,43^ and transmembrane protein 106B (TMEM106B)^5,9,18,20,24,28,36,44^. Furthermore, a component of similar morphology, albeit in non-lysine environments, was found in amyotrophic lateral sclerosis with frontotemporal lobar degeneration (ALS-FTLD)^2^, FTLD-TDP^3^ and aFTLD^36^, involving the misfolded proteins TAR DNA-binding protein 43 (TDP-43)^2,3^ and TATA-binding protein-associated factor 15 (TAF15)^36^.

Neuropathological fibrils comprise multiple protein molecules arranged in a stack, like a pile of plates. The individual protein molecules are misfolded flat, rather than having the more usual globular shape of proteins. The stacking distance is about 4.8 Å in all fibrils regardless of disease and protein type. Typically, LCEDs are elliptic rods, running parallel to the fibril and at right angles to the protein, consistent with a structural role as guide and support (Fig. 1). LCEDs are continuous, with detailed features and a repeat distance matching that of protein, consistent with a polymer (Fig. 2). LCEDs from completely different diseases, involving completely different proteins, have similar features and connectivity, pointing to a common constituent molecule (Fig. 3).

**Fig. 1.**
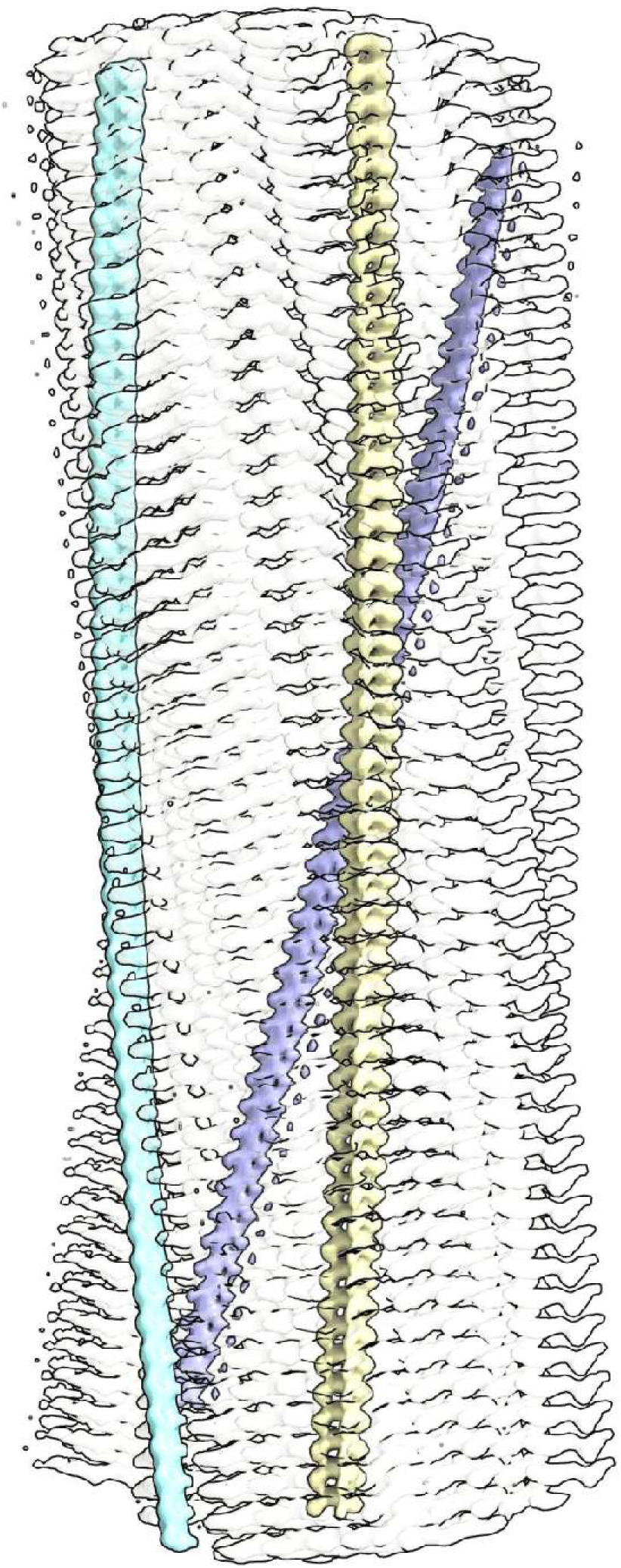
Lysine coordinating extra densities are like supporting rods. Alpha-synuclein fibril of multiple system atrophy. LCEDs (colours) are upright, parallel to the fibril and at right angles to protein rungs (transparent, white), consistent with a structural role as guide and support. At authors’ contour level. Distance between protein rungs is 4.7 Å. Image of EMD-10650^27^ created with UCSF ChimeraX^46^.

**Fig. 2.**
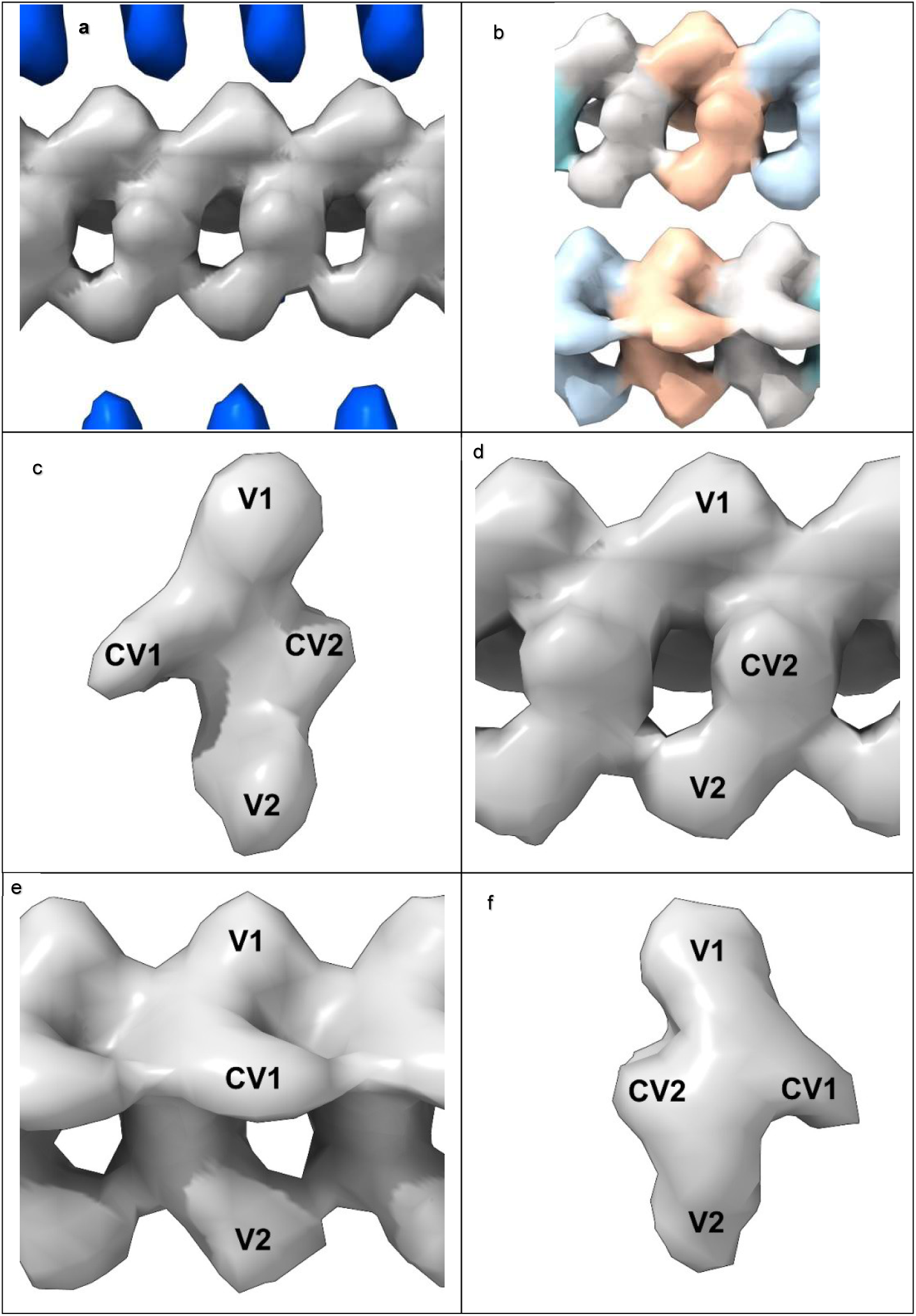
LCEDs are consistent with an underlying polymer. LCED at K43 in α-syn fibril of MSA, at authors’ contour level. **a.** LCED (grey) is at right angles to the protein (blue rungs). **b.** Repeat distance 4.7 Å. **c.-f.** Elliptic rod with distinct features at vertices, V1 and V2, and co-vertices, CV1 and CV2. The well-defined repeating features, continuous and in step with protein, are consistent with an underlying polymer. Images of EMD-10650^27^ created with UCSF ChimeraX^46^. Colour-shading of repeating units in **b.** by Segger^64^.

**Fig. 3.**
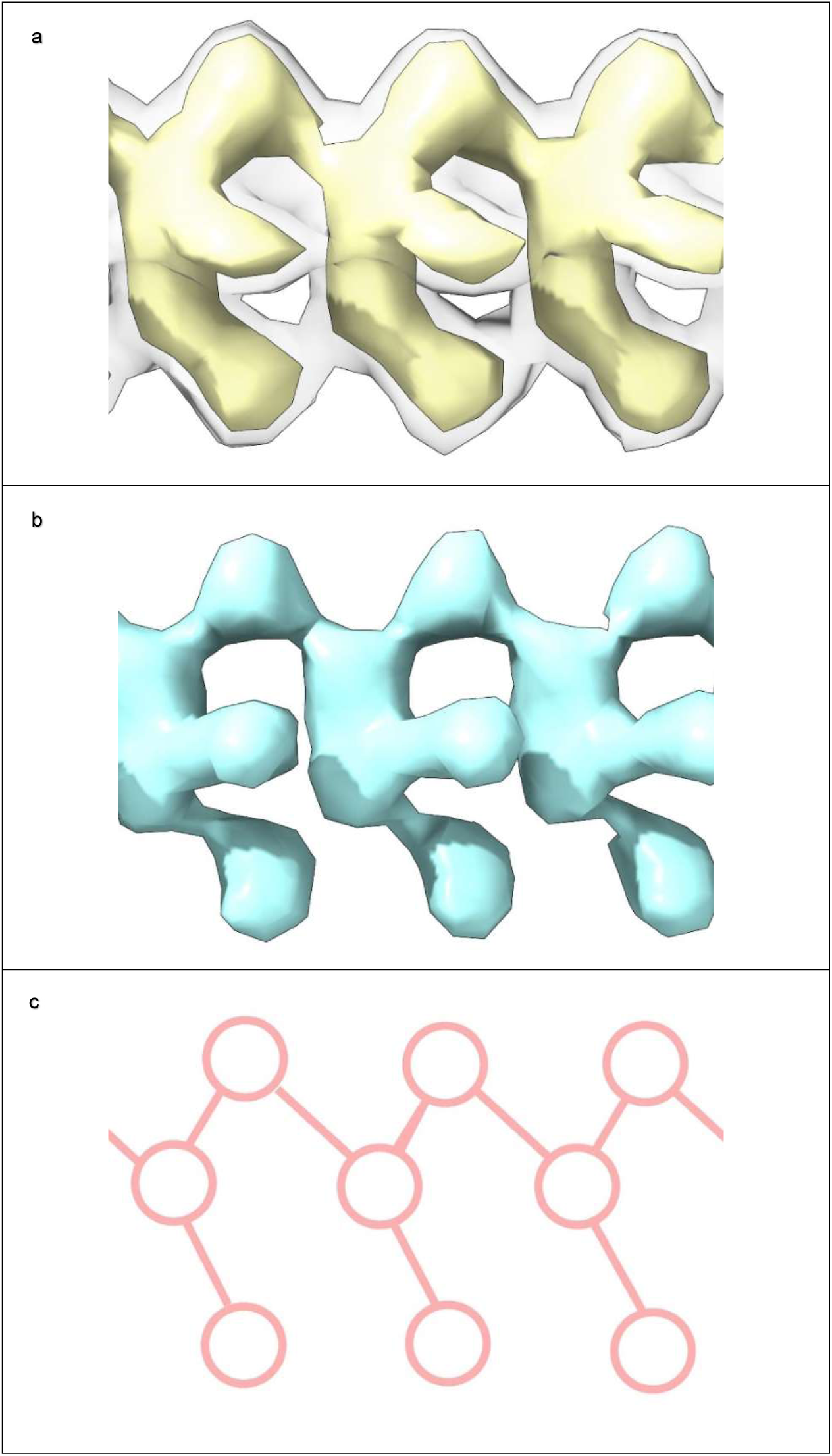
LCEDs are consistent with a molecular factor in common. **a.** LCED from α-syn fibril of MSA, at two contour levels (authors’ level transparent grey). **b.** tau fibril of LNT, at authors’ level. The LCEDs, involving completely different proteins and diseases, have common features and Y-shaped connectivity, consistent with a constituent molecule in common, a polymer. Images of EMD-10650^27^ and EMD-13225^33^ created with UCSF ChimeraX^46^. Connectivity diagram (**c**) created with ImageJ^65^.

LCEDs were graded as weak, moderate or strong (see Methods). This property of strength reflects the occupancy relative to protein. In paired helical filaments (PHFs), at the K317-K321 locus, LCEDs are present in 66.7% of assessable maps, and, where present, vary in strength from weak (29.7%), to moderate (12.5%) to strong (57.8%). This suggests a dynamic relationship, whereby the constituent molecule binds and unbinds to protein and feasibly passes between fibrils. It does not negate a structural role, since the constituent molecule could act as a temporary rather than permanent scaffold for fibril construction in some instances.

In 82% of cases LCEDs are elliptic rods (Extended Data Fig. 1) with either a Y-shaped (51.1% of LCEDs, Extended Data Fig. 2) or X-shaped substructure (30.9% of LCEDs, Extended Data Fig. 3). Although, at first sight, this might suggest different constituent molecules, another interpretation is possible: the X-shaped substructure could be due to averaging together of alternative poses of the Y-shaped substructure, arising from alternative binding poses (Extended Data Fig. 3) or 2-fold symmetry (Extended Data Fig. 4i). This is in the same way as a forward-slanting Y, if superimposed on a backwards-facing Y, or lambda, or reverse lambda, would form an X.

In 12% of cases, the LCED is bulky, consistent with a duplex (Extended Data Fig. 5). In 1.3% of cases, the substructure is V-shaped, consistent with a truncated form of the Y-shaped substructure. In a single case, there is 3-fold symmetry (Extended Data Fig.4f). In only 4.3 % of LCEDs is the morphology indeterminate. These morphologies are seen regardless of strength, although in weaker densities they should be viewed at reduced contour level.

## The protein environment

In 85% of cases, LCEDs are within 3.5 Å of lysine terminal nitrogens or other coordinating atoms. At 4.8 Å, the figure is 99%. This suggests that, allowing for a degree of uncertainty about the occupancy of LCEDs and the position of atoms in the matching atomic models, protein and the constituent molecule are within hydrogen bonding distance and feasibly form hydrogen bonds and other intermolecular bonds such as van der Waals interactions. A convenient way of categorizing protein environments is by their potential for hydrogen bonding (from either purely donor or mixed donor and acceptor atoms) and the number of coordinating residues, 1-6 (see Extended Data Table 1 and Extended Data Fig. 4).

Lysine is the pre-eminent coordinating residue, but other basic residues (arginine and histidine), polar neutral residues (asparagine, glutamine, serine, threonine and tyrosine) and acidic residues (aspartate and glutamate, albeit only rarely) may also take part. A common motif is paired lysines, seen in tau and α-syn fibrils. Where LCEDs occur between two protofilaments, as in tau straight filaments (SFs) in AD and α-syn MSA fibrils (Extended Data Figs. 4h, k, l and 6a) they may act as an intermolecular glue.

As shown (Fig. 2), a typical LCED has distinctive features at its vertices and co-vertices and is therefore chiral. Hence it is possible to assign a notional elevation and direction relative to its protein environment. For example, some LCEDs are horizontal and others vertical relative to a given lysine pair (Extended Data Fig. 6b and c). Interestingly, the pose of LCEDs may vary in a given protein environment, consistent with alternative binding poses of the constituent molecule. For example, LCEDs in two different preparations of tau fibrils from LNT have the same vertical alignment but one is upside down relative to the other (Extended Data Fig. 7). Possibly, a range of presentations and variants of the constituent molecule are permitted at a particular site in the protein, where binding may be competitive or preferential rather than exclusive.

Previous authors have inferred that the constituent molecule is likely anionic^5,7,11,12,17,27,30,37,39,41,43,51^. The present analysis suggests that the situation may be more nuanced, and whilst largely anionic, as evidenced by the pre-eminence of lysines in protein environments, parts of the constituent molecule may be cationic, as evidenced by a coordinating role, in some instances, for neutral and acidic residues. For example, in the AGD fold, a glutamate, E372, coordinates with the LCED along with three lysines, K290, K294 and K370 (Extended Data Fig. 6e). In the rather similar CBD fold, only the three lysines coordinate with the LCED (Extended Data Fig. 6d). This suggests a subtle difference in the constituent molecule in the two diseases, with a consequent effect on the specific folding of the protein.

## EDs at non-lysine environments (non-LCEDs)

Rod-like densities with Y-shaped substructure, similar to LCEDs were present at Q303, N306 and Q343 in TDP-43 fibrils and at Q60 in TAF15 fibrils (Extended Data Fig. 8). Whilst these environments differ in charge to those of LCEDs, and might therefore indicate different disease mechanisms^2^, they do in fact offer similar opportunities for hydrogen bonding, taking into account the adaptive possibilities of neutral polar residues towards a potential ligand. Furthermore, the backbone atoms of glycine might conceivably be available for hydrogen bonding, depending on the geometrical relationship with the ligand, and/or there may be a role for van der Waals interactions^2^.

Notable, also, are bulky non-LCEDs at Y10 in Aβ fibrils from patients with AD and cerebral amyloid angiopathy. In one such preparation, EMD-18508, extra densities at Y10 are bulky but singleton. In the other preparation, EMD-18509, from the same paper^42^, extra densities at Y10 explicitly form a duplex of rods, in which marker placement suggests a duplex of Y-shaped chains running in anti-parallel directions (Extended Data Fig. 5).

## The constituent molecule

It has previously been asserted, on the basis of evidence in vitro, that the continuous appearance of extra densities might arise from averaging of ions out of step with the helical rise^51^. However, this would predict a blurring of features, and is therefore at variance with the evidence here concerning LCEDs from human brains ex vivo. Only 4.3% of LCEDS were deemed to be morphologically indeterminate in the present study. The remainder showed well-defined features clearly in step with the helical rise (Fig. 2). The appearances here suggest a linear straight polymer with a repeat distance of about 4.8 Å matching that of protein, mainly as a single chain but in some cases as a duplex of chains. This considerably narrows the number of candidates for the constituent molecule since there are relatively few bioavailable polymers. When combined with its inferred mainly anionic nature, the chief candidates are RNA, heparan sulphate and acidic polypeptides. Similar arguments apply to scrapie fibrils in animals where it was concluded that RNA is the most likely candidate^52^.

Scrapie is a transmissible neurodegenerative disease in sheep and laboratory animals in which misfolded PrP is associated with extra densities coordinating with lysines, arginines and histidines. Previous studies, with acridine orange, demonstrated RNA in neuropathological lesions from AD, ALS/PDC, CBD, Down’s, PiD and PSP, although not from DLB, MSA and PD^53,54^. Similarly, RNA was shown in PrP aggregates in hamster scrapie^55^.

An obvious argument against RNA is that it does not typically form a rod-like molecule with a 4.8 Å repeat distance. However, there is key experimental evidence in support of RNA. In experiments carried out in the Eisenberg lab^56^, tau was incubated with RNA and the resulting fibrils were viewed with cryo-EM. In addition to stacks of tau protein, extra densities were found, rod-like, axially continuous, at right angles to protein, within hydrogen bonding distance of arginines and histidines, with a Y-shaped substructure similar to that of LCEDs from fibrils ex vivo from human brains described here (Extended Data Fig. 9).

The atomic model of the straight form of RNA provided by the Eisenberg lab^56^ was analysed as part of the present study, by submitting it to the MolProbity website^57^. It has an unknown rotamer engendering a straight polymer but otherwise perfect geometry with normal bond lengths, normal bond angles, normal sugar puckers and a MolProbity clash score of zero. This in vitro fibril, a complex of RNA and tau, thus incorporates a straight form of RNA with a 4.8 Å repeat distance matching that of protein, indicating the feasibility of such a component in other neuropathological fibrils. Perhaps such a novel rotameric form of RNA is only found in complexation with protein, explaining why it has not previously been described. Interestingly, a straight form of nucleic acid was postulated by Watson and Crick^58^ and was also modelled in silico in α-syn MSA fibrils, where the term ortho-RNA (oRNA) was used^59^. The feasibility of RNA as the constituent molecule of LCEDs is also supported by molecular docking studies^52,59^

The authors of the Eisenberg model of RNA^56^ attributed the 3 blobs in the density to phosphate, ribose and base. A 3-blob, Y-shaped substructure is also a major structural motif of LCEDs in the present study. It is also seen in rod-like densities in non-lysine environments in TAF15 and TDP-43 fibrils.

Due to averaging, it is not possible to read a polymer sequence in these maps but, since the protein interface is repetitive, it is possible that sequence variants of the putative common polymer evidenced here are also repetitive. Host-like repetitive nucleic acid sequences have been described in brains from hamsters with scrapie^60,61^.

The evidence here of an extra density containing two anti-parallel chains (Extended Data Fig. 5e and f) provides additional evidence in support of RNA. Although the structure is not typical of double stranded RNA it could conceivably represent a refolded form of dsRNA. A similar example of an anti-parallel duplex was found in the animal disease scrapie^52^.

## Unitary basis of neurodegeneration

Evidence here suggests that a common molecular factor, a predominantly anionic polymer, is present in neuropathological fibrils from the brains of patients with AD, PD, ALS and other neurodegenerations. This suggests a unitary basis for these diseases and may explain their many clinical and pathological features of overlap. It would for example provide a simple explanation for the otherwise vexed issue of co-pathologies^47,48^. In general, several misfolded proteins may be present, as major or minor players, in a given neurodegeneration. This situation of multiple protein folds in the same brain is difficult to reconcile with the popular idea that the “strain” of a neurodegenerative disease is somehow enciphered in the protein fold^62^. Whilst this idea might seem attractive where a single fold is present, the common situation where multiple folds are present gives rise to a rather awkward interpretation, of a disease being due to a mixture of strains.

The best known example of a co-pathology is Alzheimer’s disease, in which neuropathological fibrils of Aβ and tau occur in the same brain. There has been heated debate about how this occurs and which comes first, including the well-known amyloid cascade hypothesis in which it is claimed that Aβ pathology always comes first^63^. However, the idea of a non-protein common molecular factor as promoter and structural determinant for both Aβ and tau fibrils provides a simpler explanation that does not require primacy of one fibril type over the other. The difference between AD, in which both Aβ and tau fibrils occur, and PART, in which tau but not Aβ fibrils occur, could be due to sequence variants in the common polymer. Each of the various neurodegenerative diseases might be attributable to a particular sequence variant of the common polymer, in which each sequence variant has a particular propensity to co-assemble with proteins to form particular folds. Thus, being a polymer, the idea of a common molecular factor readily explains both the unity and diversity of these diseases.

This suggests a new approach to treatment. Current treatments involve immune targeting of a single misfolded protein^47,48^. However, therapeutic clearance of Aβ pathology in AD, for example, leaves tau fibrils intact and the clinical results have been disappointing. Where several co-pathologies exist a cocktail of treatments against various protein fibrils might work better. However, targeting of a common molecular factor, as implicated in the current study, might offer a simpler and more effective treatment for all of these diseases.

Also, the idea that sequence variants of a common polymer underly the diversity of these diseases suggests the possibility of simpler and more definitive diagnostic tests in the future, by sequencing.

Clearly, further work is needed, to identify the common factor. Candidate ligands (such as repetitive RNAs) and target proteins could be mixed in vitro and in silico to determine structural changes and free energies of binding, or introduced into cell lines and experimental animals, to see whether neuropathological fibrils are produced. It would behove experimenters to treat such candidate molecules as a potential biohazard. RNA sequencing, including specialist techniques for repetitive RNAs, could be carried out on brain tissues from patients with AD and other neurodegenerations.

## Methods

### Data sources and selection

Data for this study were sourced from public repositories, the EMDB^49^ and the PDB^50^. The scope of this study was to examine all available cryo-EM maps and atomic models of neuropathological fibrils from autopsy brains of patients with AD and other neurodegenerations. Search terms used were the names of diseases and proteins including Aβ, α-syn, PrP, TAF15, tau, TDP-43 and TMEM106B.

The following maps and atomic models were accessed from the EMDB and PDB. The EMDB and PDB numbers are given in matched pairs and bracketed together for each reference. Nil is stated where no atomic model is available. Human ex vivo neuropathological fibrils were as follows: (21200, 6vh7; 21201, 6vha; 21207, 6vhl^1^);(13708, 7py2^2^);(16628, 8cg3; 16642, 8cgg; 16643, 8cgh; 16677, nil; 16681, nil; 16682, nil^3^);(50621, 9fof; 50628, 9for^4^);(26268, 7u0z; 26273, 7u10; 26274, 7u11; 26275, 7u12; 26276, 7u13; 26277, 7u14; 26278, 7u15; 26279, 7u16; 26281, nil; 26282, nil; 26283, nil; 26284, nil; 26285, nil; 26286, nil; 26287, nil; 26288, nil; 26289, nil; 26290, nil; 26291, nil; 26292, nil; 26293, nil; 26294, nil; 26295, nil; 26296, nil^5^);(0077, 6gx5; 0078, nil^6^);(0259, 6hre; 0260, 6hrf^7^);(0527, 6nwp; 0528, 6nwq^8^);(33054, 7×83; 33055, 7×84^9^);(40411, 8seh; 40413, 8sei; 40416, 8sej; 40419, 8sek; 40421, 8sel^10^);(3741, 5o3l; 3742, 5o3o; 3743, 5o3t; 3744, nil^11^);(16035, 8bgs; 16039, 8bgv^12^);(29036, 8ff2; 29037, 8ff3; 29038, nil^13^);(50587, 9fnb^14^);(45005, 9pxi; 45007, 9bxo; 45008, 9bxq; 45009, 9bxr^15^);(23871, 7mkf; 23890, 7mkg; 23894, 7mkh^16^);(26607, 7umq; 26613, 7un5^17^);(25995, 7tmc; 28943, 8f9k^18^);(46417, 9czi; 46420, 9czl; 46422, 9czn; 46424, 9czp^19^);(24953, 7saq; 24954, 7sar; 24955, 7sas^20^);(42463, 8uq7^21^);(29458, 8fug^22^);(16532, 8caq; 16535, 8cax^23^);(17171, 8ot6; 17173, 8ot9; 17174, 8otc; 17175, 8otd; 17176, 8ote; 17177, 8otf; 17178, 8otg; 17179, 8oth; 17180, 8oti; 17181, 8otj^24^);(19846, 9eo7; 19849, 9eo9; 19852, 9eoe; 19854, 9eog^25^);(19926, 9erm; 19927, 9ern; 19928, 9ero^26^);(10650, 6xyo;10651, 6xyp;10652, 6xyq^27^);(14174, 7qvc;14176, 7qvf;14187, 7qwg;14188, 7qwl;14189, 7qwm^28^);(17383, 8p34^29^);(51319, 9gg0;51320, 9gg1;51325, 9gg6^30^);(26663, 7upe;26664, 7upf;26665, 7upg^31^);(12549, 7nrq;12550, 7nrs;12551, 7nrv;12552, 7nrx;12553, nil^32^);(13218, 7p65;13219, 7p66;13220, 7p67;13221, 7p68;13223, 7p6a;13224, 7p6b;13225, 7p6c;13226, 7p6d;13227, 7p6e^33^);(16329, 8byn^34^);(15770, 8azs;15771, 8azt; 15772, 8azu^35^);(16999, 8ons;17020, nil;17021, nil; 17022, nil; 17109, nil; 18226, nil; 18227, nil; 18236, nil; 18240, nil; 18241, nil; 18242, nil; 18243, nil^36^);(13800, 7q4b; 13809, 7q4m; 16434, nil^38^);(15285, 8a9l^39^);(16022, 8bfz; 16023, 8bg0^40^);(16188, 8bqv; 16189, 8bqw^41^);(18508, 8qn6; 18509, 8qn7^42^);(45464, 9cd9; 45465, 9cda; 45979, 9cx6^37^);(10512, 6tjo; 10514, 6tjx^43^);(36043, 8j7n; 36045, 8j7p; 38069, 8×5h^44^). An experimental RNA-induced tau fibril (EMD-25364, PDB 7sp1^56^) was also studied. A total of 161 cryo-EM maps were examined^1–44^, involving 7 proteins, Aβ^10,13,19,24,35,36,38,40,42^, α-syn^27,37,39,41^, PrP^17^, TAF15^36^, tau^1,5–8,10–12,15,16,19,21–26,29–35,43^, TDP-43^2–4^ and TMEM106B^5,9,14,18,20,24,28,36,44^ and over 30 neurodegenerative diseases, including Aβ-cerebral amyloid angiopathy (Aβ-CAA)^13,42^, AD^1,7,11,12,21,22,28,31,32,35,38^, AD with cotton wool plaques^19^, AD with Artic mutation E22G^40^, AD with R406W mutation^25^, AD with V337M mutation^25^, atypical frontotemporal lobar degeneration (aFTLD)^36^, argyrophilic grain disease (AGD)^33^, amyotrophic lateral sclerosis with frontotemporal lobar degeneration (ALS-FTLD)^2^, corticobasal degeneration (CBD)^1,43^, chronic traumatic encephalopathy (CTE)^8^, dementia with Lewy bodies (DLB)^5,39^, Down’s syndrome^10,15^, frontotemporal dementia and parkinsonism linked to chromosome 17 (FTDP-17) P301L^30^, FTDP-17 P301T^30^, frontotemporal lobar degeneration (FTLD-TDP) type A^3,5^, FTLD-TDP type B^5^, FTLD-TDP type C^4,5,20^, globular glial tauopathy (GGT)^33^, Gerstmann-Sträussler-Scheinker disease (GSS)^16,17^, amyotrophic lateral sclerosis/parkinsonism-dementia complex (ALS/PDC) of Guam^24^, juvenile-onset synucleinopathy (JOS)^41^, ALS/PDC of Kii^24^, limbic-predominant neuronal inclusion body 4R tauopathy (LNT)^33^, multiple system atrophy (MSA)^27,28,37^, multiple system tauopathy with presenile dementia (MSTD)^18^, primary age-related tauopathy (PART)^32,34^, posterior cortical atrophy (PCA)^32^, Parkinson’s disease (PD)^9,39^, Pick’s disease (PiD)^6,29^, prion protein-cerebral amyloid angiopathy (PrP-CAA)^16^, progressive supranuclear palsy (PSP)^5,33^, subacute sclerosing panencephalitis (SSPE)^23^ and vacuolar tauopathy (VT)^26^. Maps from Biondi bodies^14^ and normal elder(s)^9,44^ were also examined.

Lysine-coordinating extra densities (LCEDs) were the primary subject of the present study. Extra densities (EDs) from other environments (*e.g.* the hydrophobic cavity in tau fibrils from CTE^8^) were generally excluded. However, a small number of EDs, morphologically similar to LCEDs but from non-lysine environments in TAF15, TDP-43 and Aβ fibrils, were also studied, responsive to the observation^2^ that they might inform of different disease mechanisms.

EDs were named by the EMDB number and the first coordinating residue. For example, ED 10650-43 is the ED in EMD-10650 opposite residue K43 in the matching atomic model, PDB 6xyo. Where pairs of LCEDs were present in doublets, the associated protofilaments were denoted “a” and “b” (left and right respectively, in the opening position).

Of the 161 maps that were found in the EMDB^1–44^, 126 had matching atomic models in the PDB. Twenty maps were considered unassessable for LCEDs, due to lack of matching atomic models and/or map factors such as low resolution or noise. In total, 233 LCEDs were found in the 141 assessable maps. Additionally, 13 EDs from non-lysine environments were studied.

### Observations and measurements

Observations and measurements were made in UCSF ChimeraX^46^ unless otherwise stated. It is important to emphasise that the present study, using cryo-EM data, is on neuropathological material from actual patients, an important distinction, in that more traditional atomic-level structural techniques, such as X-Ray diffraction, are typically confined to *in vitro* preparations. Studies on actual human tissue are uniquely eloquent about the mechanisms and causes of diseases, notwithstanding the complementary benefits accruing from *in vitro* work^45,51^. In effect, ChimeraX acts as a microscope for atomic-level autopsy (nano-autopsy) of patients with AD and other neurodegenerations.

A Dell Desktop-GUQ6HDT XPS 15 9500 with Intel® Core™ i7-10750H CPU @2.6 GHz with 16 GB RAM running Windows 11 Home (version 21H2) with broadband internet connection was used to run software and to access data and software online.

Each cryo-EM map was examined at the authors’ recommended contour level with the protein model *in situ*. For clarity, the map was made transparent and clipped to the region of the protein. LCEDs were identified in their protein environment, by inspection and reference to the published papers. Allowing for differences in resolution and occupancy, LCEDs from the subject group were similar (*e.g.* they were straight, continuous, repeating and parallel to the fibril axis).

To determine whether they were in perfect register with the protein, markers were placed 30 units apart on LCEDs and 30 rungs apart on the adjacent protein density and the distances compared. To determine whether LCEDs maintained a constant distance and attitude to the protein, slices of the fibril, 5 Å in thickness and 100 Å apart, were compared with the fitmap command. To assess whether LCEDs were within hydrogen-bonding distance of protein, color zones from the NZ (terminal nitrogen) atoms of lysines or equivalent atoms in other coordinating residues were incremented until they reached the surface of the LCEDs.

### Nanodissection of LCEDs

In well-resolved cases, LCEDs were detached from, or showed only minor regions of fusion to, the protein density. Some EDs, generally at lower resolution, were more extensively fused to the protein density. A technique was developed to dissect out LCEDs from the rest of the density, at nanoscale level, using the map eraser tool, in order to examine them unobscured by the protein density. The LCED was erased and the remainder subtracted from a copy of the original.

ED 10650-43^27^ was chosen as a reference LCED, for comparison with other LCEDs, as it has relatively high resolution (2.6 Å) and strong occupancy. It is possible that the exquisite detail seen in this LCED is also due to it being within a cavity and strongly braced by five coordinating protein residues at each rung. It is an elliptic rod with distinctive features at the vertices and co-vertices (Fig. 2c-f) and resembles a series of connected Ys, slanting either forwards or backwards, depending on the side of view (Figs. 2 and 3a). Further detail was evinced by adjusting the contour level. For example, in Fig. 3a, the LCED is shown at two contour levels. At higher contour level (reduced volume), the connectivity of the LCED is more readily apparent. Using such cues, LCEDs were oriented and compared to the reference LCED. Detection and colour-shading of repeating units (Fig. 2b) was performed with Segger^64^ in UCSF ChimeraX^46^.

### Contour mapping of LCEDs

A technique of contour mapping was developed to determine the most persistent regions of density and connections within LCEDs. Copies of the cryo-EM map were viewed at different contour levels, always including the authors’ recommended level, and the outlines were superimposed, similar to a contour map of a geographical terrain.

Similarly, connections between LCEDs and protein density were determined (Extended Data Fig. 6a). Such connections were named tethering patterns and are consistent with intermolecular bonds such as hydrogen bonds. By reference to the atomic model, it was noted which protein residues formed part of the tethering pattern, whether the LCED was vertical or horizontal with respect to the protein motif (Extended Data Fig. 6b and c) and whether the LCED was on the outside of the fibril or within a groove, cleft or cavity.

LCEDs were assessed as having high (or strong, or near-stoichiometric) occupancy where, at the authors’ contour level, the volume was comparable to adjacent protein density. LCEDs were assessed as having low (or weak) occupancy where the volume was attenuated in comparison to the adjacent protein density. In order for the assessment to be strictly reproducible, the following criteria were adopted. At authors’ level, strong LCEDs were joined continuously throughout. LCEDs of moderate strength were joined at a minimum of two consecutive levels. Weak LCEDs were discontinuous throughout. For weak densities, to distinguish them from noise, the repeating units were defined as bean- or gravel-like (rather than merely dust-like) and substantially uniform in size and shape opposite each protein rung.

### Marker placement

Using the marker tool in UCSF ChimeraX^46^, markers were placed in three dimensions within LCEDs in order to deconstruct their features and connections. Once done, it was also possible to determine the specific binding poses of LCEDs relative to protein (Extended Data Fig. 7). Markers were placed in regions of major density at the vertices and middle of the elliptic outline, and links between markers were placed in regions of persistent density. The decision about placement was made considering the appearances at the authors’ contour level and other nearby levels, higher and lower.

In LCEDs with Y-shaped substructures, there were hump-like densities at one vertex and peg-like densities at the other (Extended Data Fig. 2). The markers in the hump-like densities were joined with the middle markers, to form a zig-zagging backbone, each marker joined with two others. On the other hand, markers in the peg-like densities at the other vertex, were joined just once, with a middle marker, and thus appeared as side-groups coming off the backbone. Either at authors’ contour level, or an adjacent level, it was apparent that only one possible join was permitted for markers in the peg-like densities.

In LCEDs with X-shaped substructure (Extended Data Fig. 3b, d and f), there was a double-hump appearance, with symmetrical hump-like densities at both vertices of the elliptic outline. The markers in the humplike densities were joined with the middle markers, each marker joined with two others. In X-shaped substructures, the density permitted different attitudes of Y (forwards-slanting Y, backwards-slanting Y, normal lambda and reverse lambda).

### Examination of other fibrils

As mentioned above, apart from LCEDs, a total of 13 rod-like densities from Aβ, TAF15 and TDP-43 fibrils at non-lysine environments, with a resemblance to LCEDs, were also examined, by similar methods, and referred to here as non-LCEDs.

In addition to fibrils from human brains, for pertinent comparison, an experimental model of RNA-induced tau fibrils^56^ (represented as EMD-25364 and PDB 7sp1) was examined in UCSF ChimeraX^46^ with similar techniques. Hydrogen bonds were displayed with relaxed criteria (distance tolerance 0.4 Å, angle tolerance 20°). The authors’ RNA model^56^, possessing an unusual rod-like structure reminiscent of the rod-like structures seen in the fibrils from human brains, was submitted to the MolProbity Webserver^57^ to determine the rotamer type, bond lengths, bond angles, sugar puckers and clashscore.

### Ethical statement

Regarding patient data, the cryo-EM maps and atomic models used in the present study were sourced from the EMDB and PDB public repositories, from the publications recorded herein, which all affirm that ethical review and informed consent were obtained and provide the names of the relevant organizations. All patient data for the present study was at one remove and no new patient data was obtained.

### Data availability

The EMDB and PDB accession numbers for publicly available data examined in this paper are provided in the original papers. They are also listed in the Methods. Any other relevant data are available from the corresponding author upon request.

## Acknowledgements

The present work was done on cryo-EM and atomic models from the EMDB and PDB public repositories. The patients and relatives were thanked in the original publications and, at one remove, I also thank them. I also thank the scientists, software developers, data managers and funders whose efforts, at one remove, have made my own work possible. Molecular graphics and analyses performed with UCSF ChimeraX were developed by the Resource for Biocomputing, Visualization, and Informatics at the University of California, San Francisco, with support from National Institutes of Health R01-GM129325 and the Office of Cyber Infrastructure and Computational Biology, National Institute of Allergy and Infectious Diseases. I thank St George’s University Hospitals NHS Foundation Trust NHS and South West London Pathology for my employment and St George’s University of London for my honorary academic status.

## Author contribution

LRB performed the whole of this work.

## Competing interests

The author declares no competing interests.

---

**Extended Data Fig. 1.**
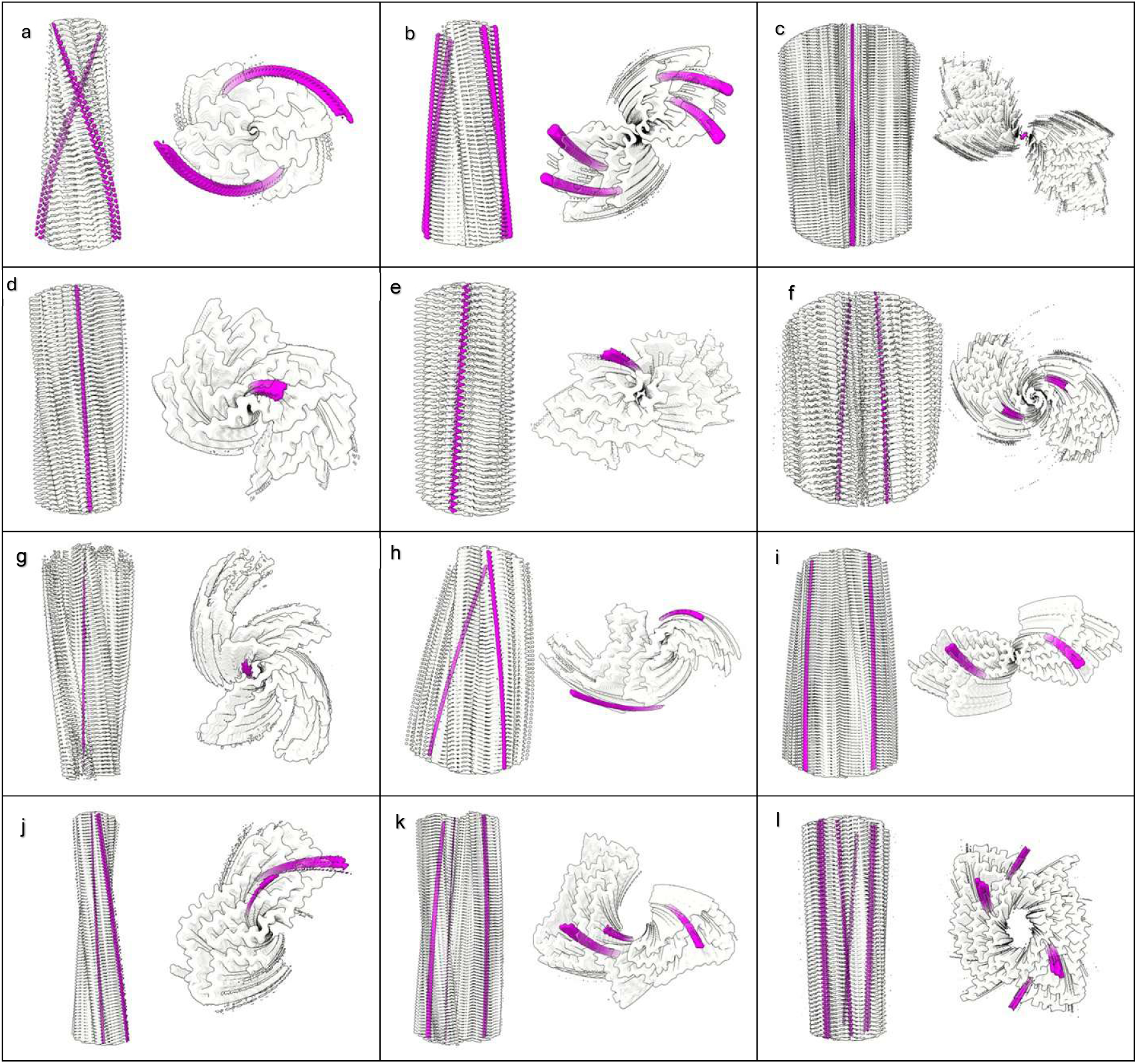
LCEDs are a common feature of diverse neuropathological fibrils. Protein white, LCEDs fuchsia, spacing of protein rungs about 4.8 Å. Images of fibrils created with UCSF ChimeraX^46^. **a.** Aβ fibril in AD, EMD-15770^35^, **b.** PrP fibril in GSS, EMD-26607^17^, **c.** TMEM106B fibril in FTLD-TDP, EMD-26290^5^, **d.** α-syn fibril in MSA, EMD-10652^27^, **e.** α-syn fibril in PD, EMD-15285^39^, **f.** α-syn fibril in JOS, EMD-16189^41^, **g.** tau fibril in AD, SF, EMD-0260^7^, **h.** tau fibril in CTE, type II, EMD-0528^8^, **i.** tau fibril in CBD, EMD-10514^43^, **j.** tau fibril in PSP, EMD-13218^33^, **k.** tau fibril in GGT, EMD-13221^33^, **l.** tau fibril in LNT, EMD-13225^33^.

**Extended Data Fig. 2.**
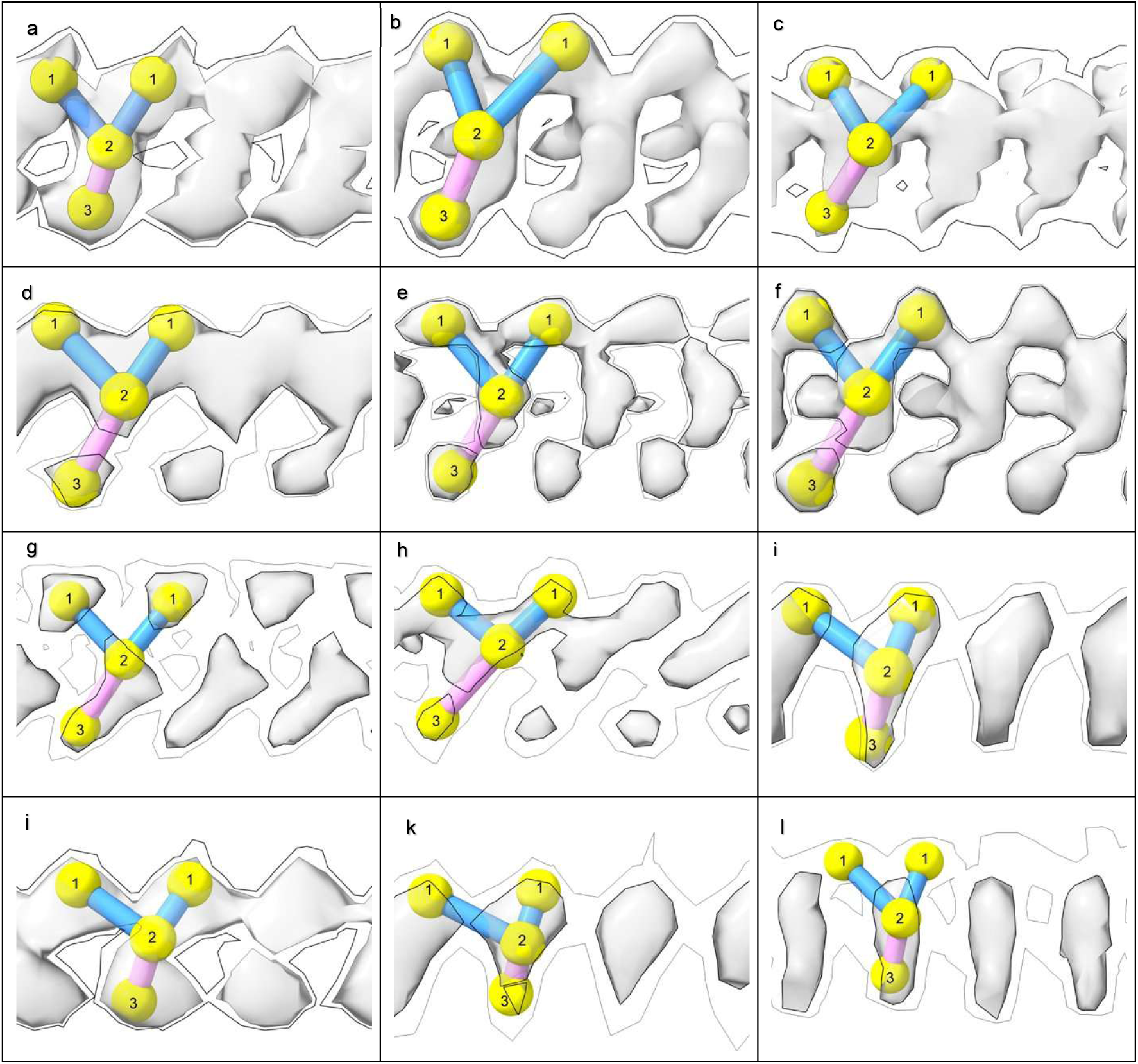
LCEDs are typically Y-shaped. Shown at two contour levels with markers, authors’ level shown with bold outline, features repeat at about 4.8 Å. Images of fibrils created with UCSF ChimeraX^46^. **a.** tau fibril in CBD at K290, EMD-10514^43^, **b.** α-syn fibril in MSA at K43, EMD-10650^27^, **c.** tau fibril in PART at K317, EMD-12550^32^, **d.** tau fibril in GGT at K317, EMD-13219^33^, **e.** tau fibril in LNT at K317, EMD-13224^33^, **f.** tau fibril in LNT at K317, EMD-13225^33^, **g.** Aβ fibril in AD at S26, EMD-15770^35^, **h.** TMEM106B fibril in aFTLD at K178, EMD-18240^36^, **i.** tau fibril in PSP at K317, EMD-26268^5^, **j.** PrP fibril in GSS at K104, EMD-26613^17^, **k.** tau fibril in AD at K311, EMD-26665^31^, **l.** TMEM106B fibril in MSTD at K139, EMD-28943^18^.

**Extended Data Fig. 3.**
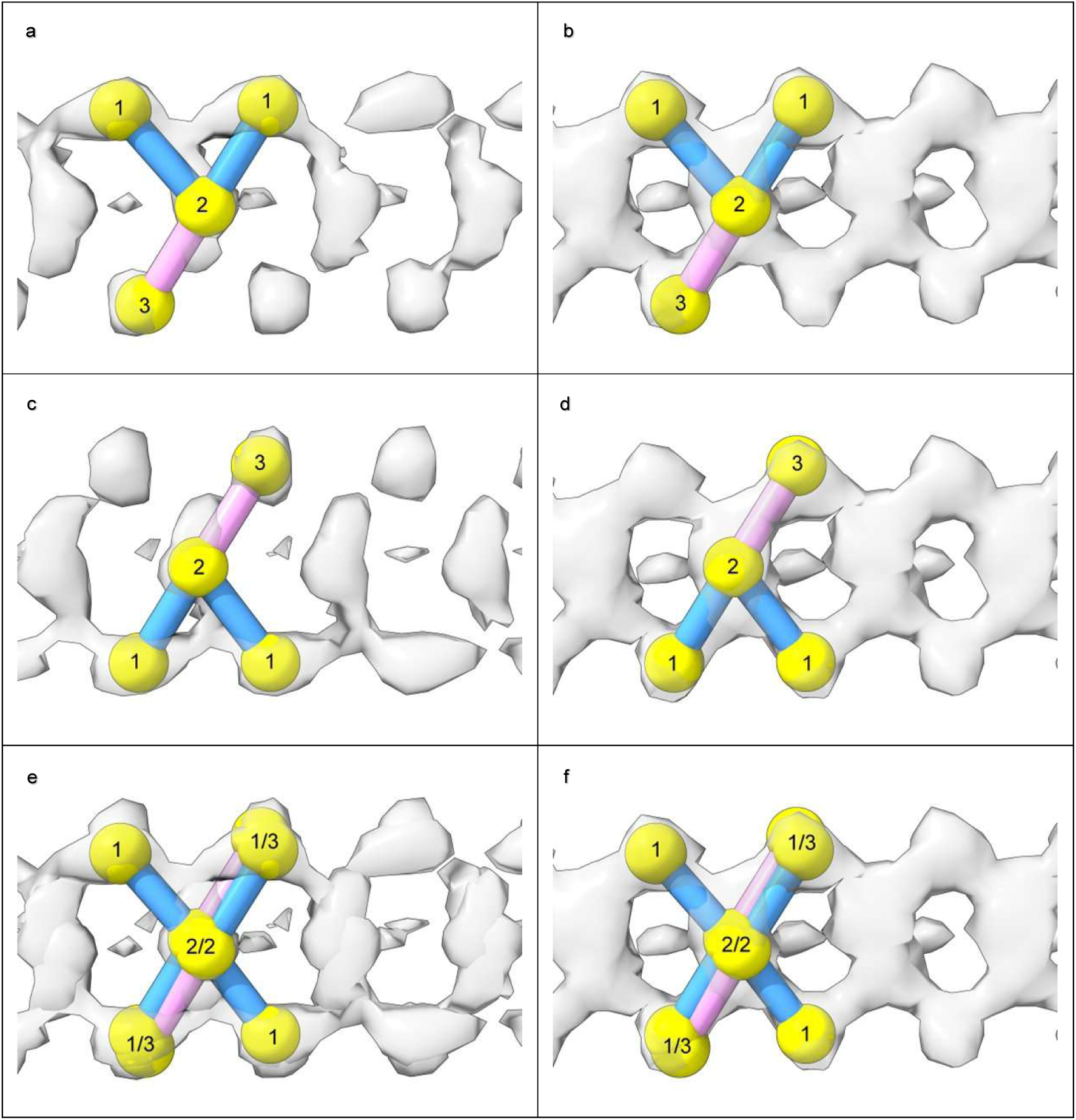
LCEDs with an X-shaped substructure are consistent with a Y-shaped constituent molecule. LCEDs shown at authors’ level with markers. Images of fibrils created with UCSF ChimeraX^46^. **a. c.** and **e.** tau fibril in LNT at K317, EMD-13224, PDB 7p6b^33^. **b. d.** and **f.** tau fibril in LNT at K317, EMD-13223, PDB 7p6a^33^. In **a.** and **c.**, viewed from different angles, the LCED is naturally Y-shaped. An X-shape was created artificially in **e.** by superimposition of the Y-shaped poses in **a.** and **c**. In **b. d.** and **f.** the LCED is naturally X-shaped, but Y-shaped markers can be fitted either way up, indicating that the X-shape is consistent with averaging of alternative Y-shaped poses.

**Extended Data Fig. 4.**
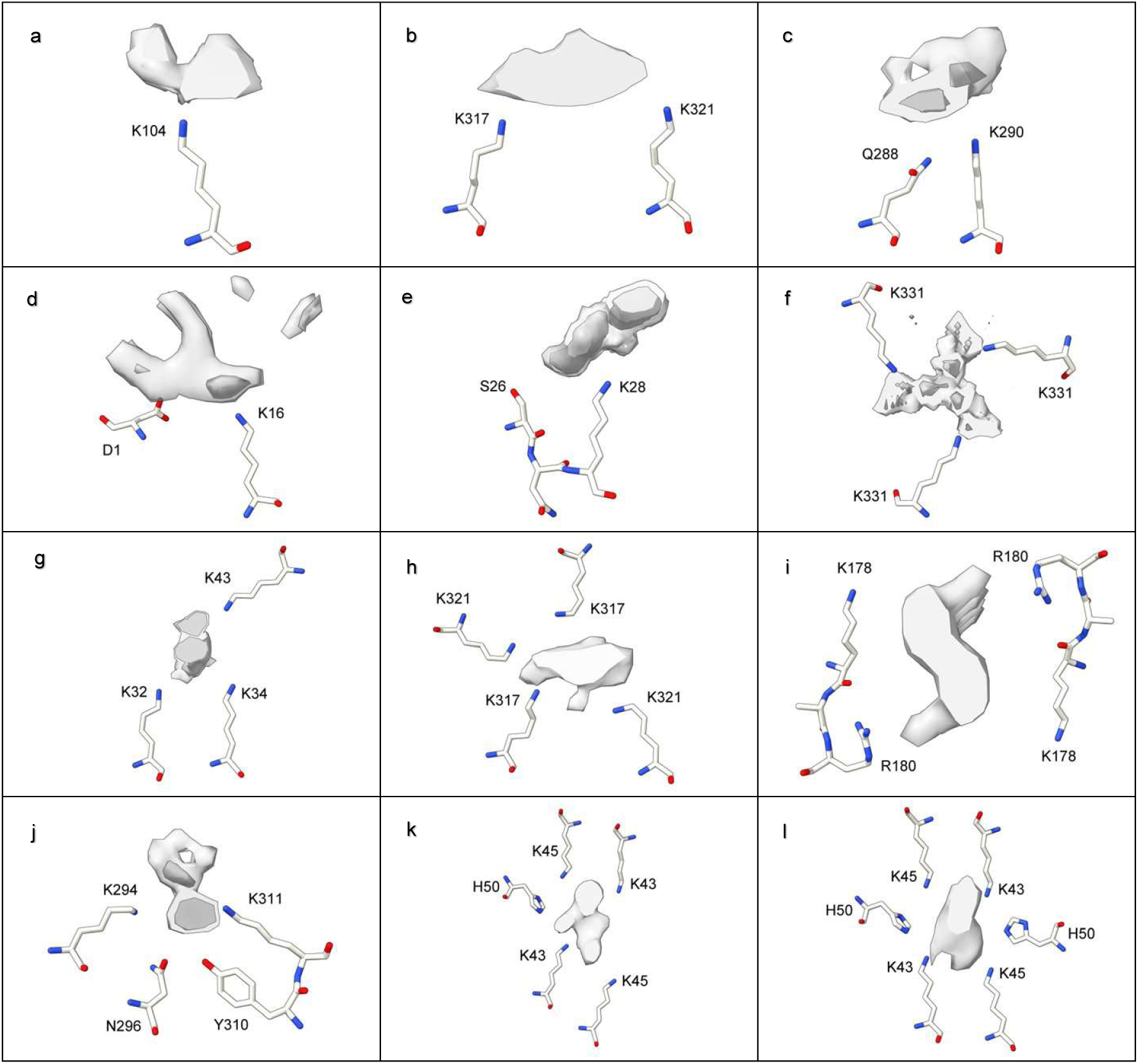
LCEDs occur in stereotyped protein environments. LCEDs in transverse section opposite 1-6 polar residues. The chemical environments are either purely donor or mixed, donor and acceptor, for hydrogen-bonding purposes. LCEDs shown at authors’ level (light grey) or at two levels (authors’ level darker and opaque). Images of fibrils created with UCSF ChimeraX^46^. **a.** PrP fibril in GSS at K104, EMD-26613, PDB 7un5^17^, **b.** tau fibril in AD, PHF, at K317, EMD-15772, PDB 8azu^35^, **c.** tau fibril in GGT at Q288, EMD-13221, PDB 7p68^33^, **d.** Aβ fibril in Down’s at D1, EMD-40421, PDB 8sel^10^, **e.** Aβ fibril in AD at S26, EMD-15770, PDB 8azs^35^, **f.** tau fibril in AD V337M, triplet filament, at K331, note 3-fold symmetry, EMD-19852, PDB 9eoe^25^, **g.** α-syn fibril in PD at K32, EMD-15285, PDB 8a9l^39^, **h.** tau fibril in PART, SF, at K317, EMD-12550, PDB 7nrs^32^, **i.** TMEM106B fibril in MSTD at K178, note 2-fold symmetry, EMD-28943, PDB 8f9k^18^, **j.** tau fibril in FTDP-17 P301T at K294, EMD-51320, PDB 9gg1^30^, **k.** α-syn fibril in MSA at K43, EMD-10650, PDB 6xyo^27^, **l.** α-syn fibril in MSA at K43, EMD-10652, PDB 6xyq^27^.

**Extended Data Fig. 5.**
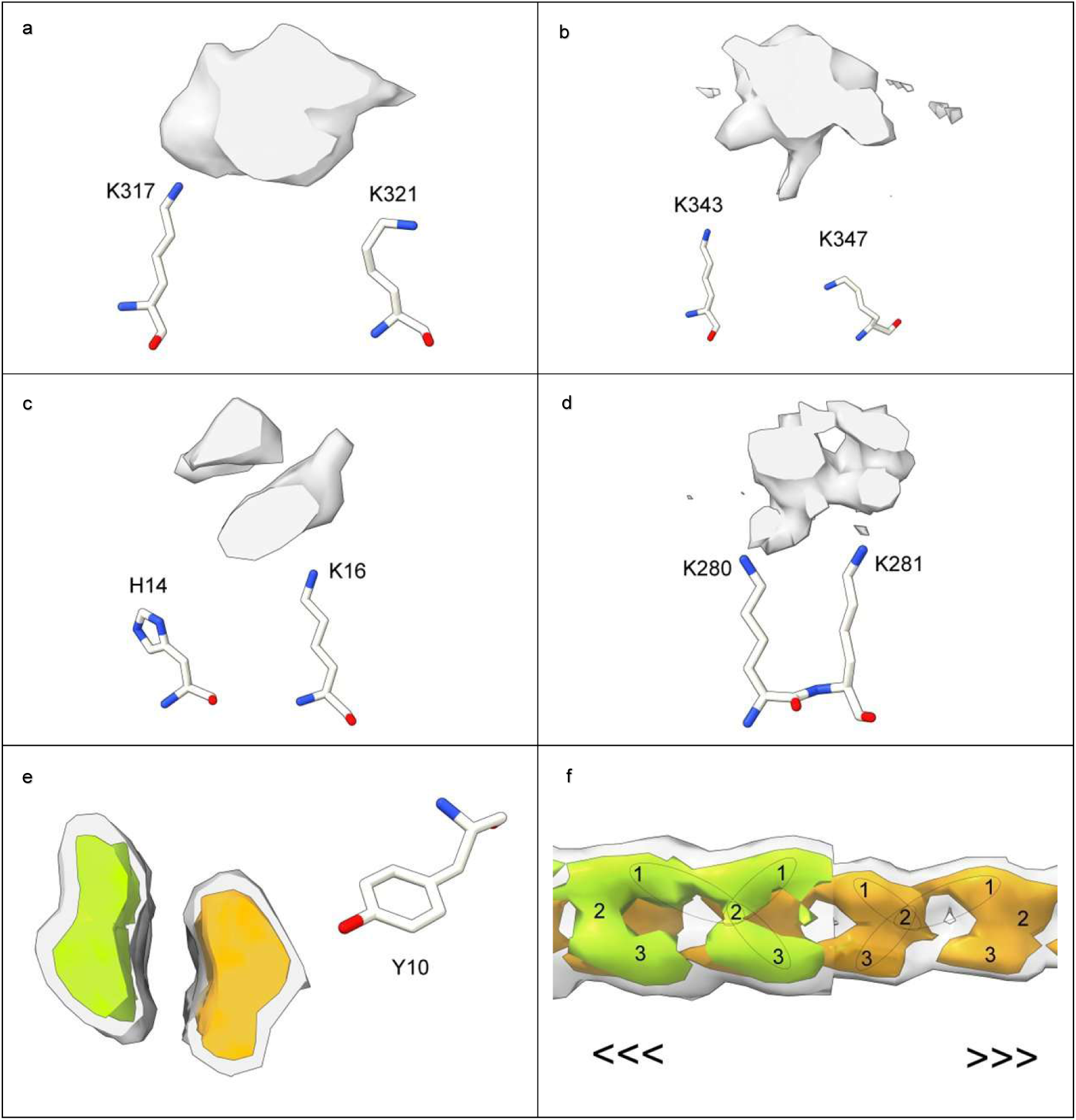
LCEDs are sometimes bulky. LCEDs in transverse section opposite protein residues at authors’ level (**a**-**d**). Also shown is a bulky non-LCED (**e** and **f**). Images of fibrils created with UCSF ChimeraX^46^. **a.** tau fibril in AD, PHF, at K317, EMD-3742, PDB 5o3o^11^. **b.** tau fibril in CBD at K343, EMD-10512, PDB 6tjo^43^. **c.** Aβ fibril in Down’s at H14, note duplex, EMD-40419, PDB 8sek^10^. **d.** tau fibril in LNT at K280, EMD-13223, PDB 7p6a^33^. **e.** and **f.** Aβ fibril in AD/CAA at Y10, note duplex, EMD-18509, PDB 8qn7^42^, authors’ level grey transparent, increased level colours opaque, in transverse section (**e**) and lengthwise (**f**). The non-LCED is explicitly divided into two chains. In **f.** one of the chains is cut away to show the other chain behind. The directions of the two chains are opposite (antiparallel).

**Extended Data Fig. 6.**
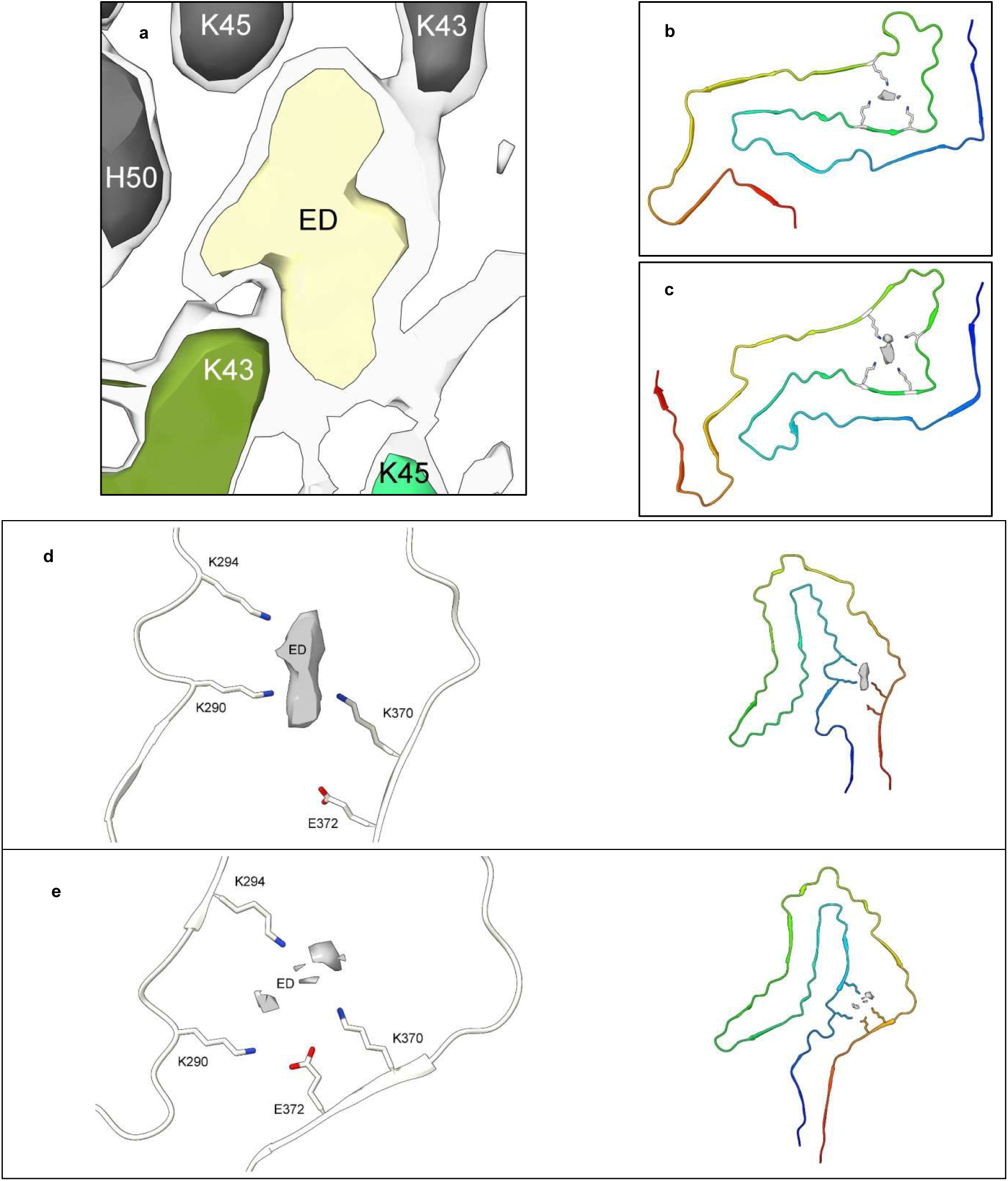
LCEDs and protein folding. The spatial relationships of LCEDs are consistent with an effect on protein folding. **a.** LCED in α-syn fibril of MSA at K43, at two levels, authors’ level opaque, note connections between extra density and protein, EMD-10650^27^. LCEDs (authors’ levels) and protein models from: **b.** tau fibril of PSP at K317, LCED lies horizontally, EMD-13218 and PDB 7p65^33^, **c.** tau fibril of GGT at K317, LCED lies vertically, EMD-13219 and PDB 7p66^33^, **d.** tau fibril of CBD at K290, EMD-10514 and PDB 6tjx^43^, **e.** tau fibril of AGD at K290, note involvement of E372 (compare to **d**), EMD-13226 and PDB 7p6d^33^. Images created with UCSF ChimeraX^46^.

**Extended Data Fig. 7.**
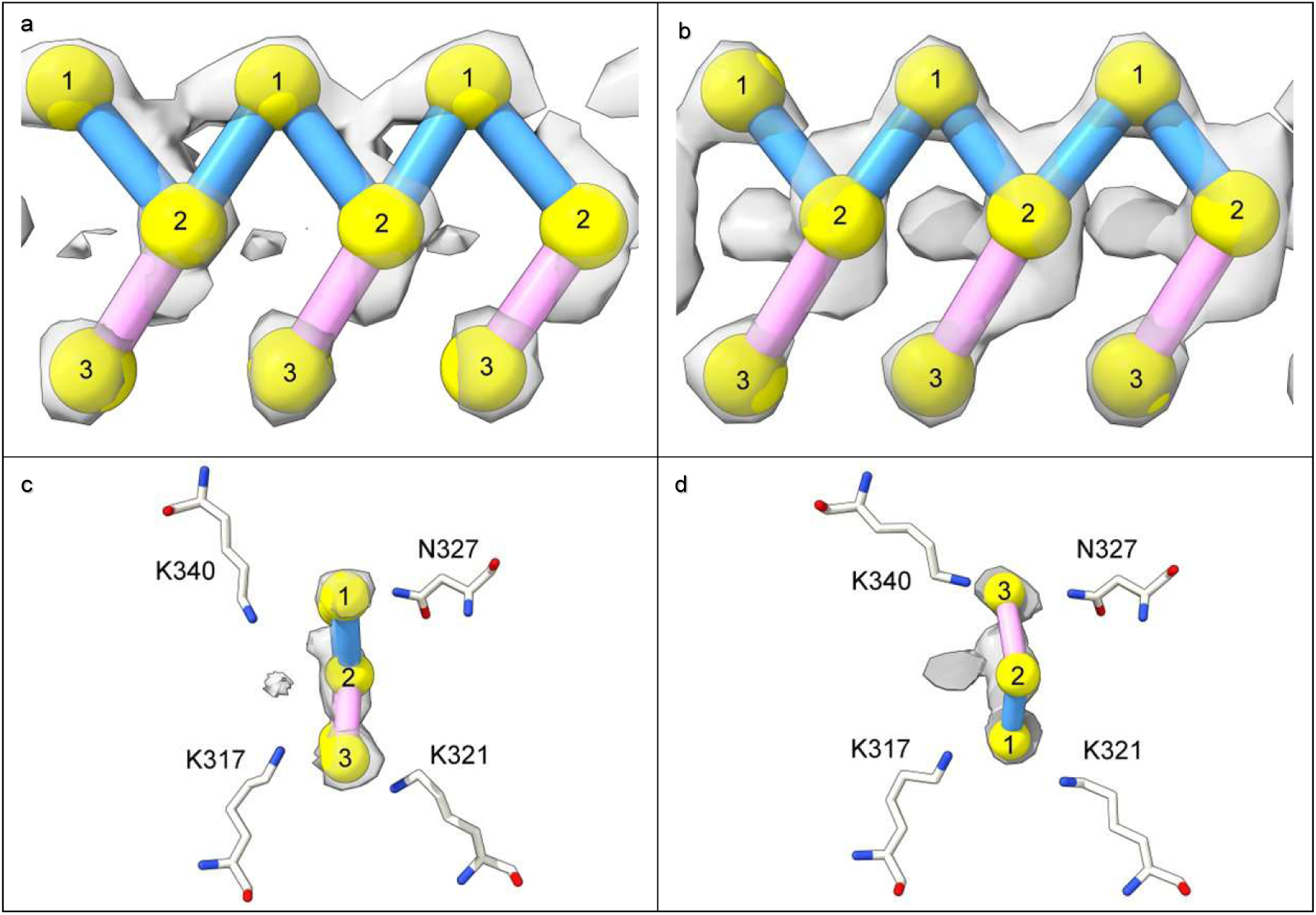
LCEDs adopt alternative binding poses. LCEDs shown at authors’ level with markers. Images of fibrils created with UCSF ChimeraX^46^. **a.** and **c.** tau fibril in LNT at K317, EMD-13224, PDB 7p6b^33^. **b.** and **d.** tau fibril in LNT at K317, EMD-13225, PDB 7p6c^33^. At top, LCEDs are shown lengthwise and at bottom they are shown in cross-section, in their protein environments. Although the morphologies are virtually identical, the binding poses are opposite, as demonstrated by the position of the markers relative to the coordinating residues in the lower frames.

**Extended Data Fig. 8.**
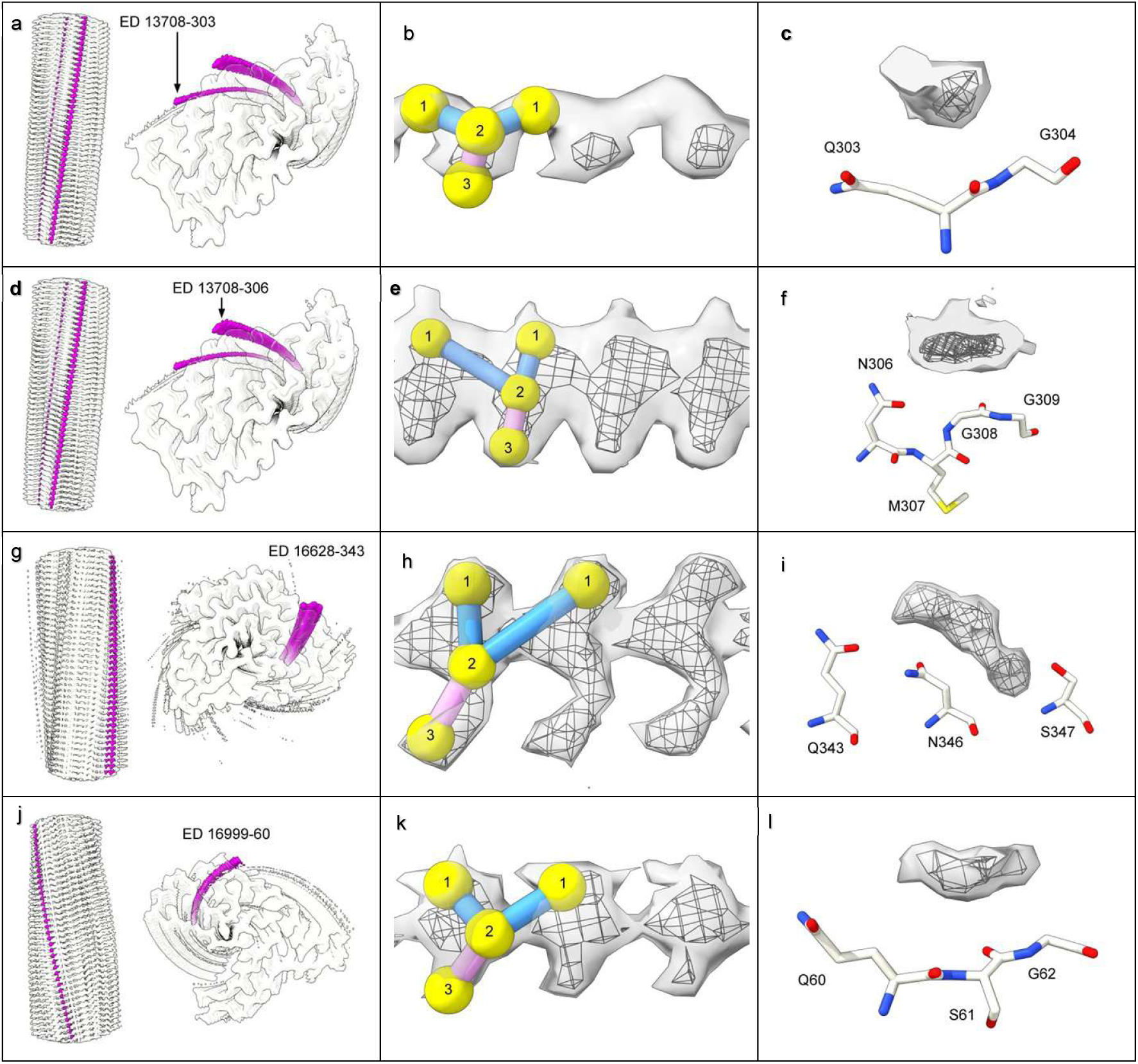
Rod-like densities also occur in non-lysine environments. Note similarities with LCEDs, including rod-like structure (**a**, **d**, **g** and **j**, protein white, non-LCEDs fuchsia, spacing of protein rungs about 4.8 Å), Y-shaped substructure (**b**, **e**, **h** and **k**, at two levels with markers, authors’ level in mesh, features repeat at about 4.8 Å) and proximity to proton donor and acceptor atoms in the protein (**c**, **f**, **i** and **l**, at two levels, author’s level in mesh, opposite coordinating residues). Images of fibrils created with UCSF ChimeraX^46^ **a. b.** and **c.** TDP-43 fibril in ALS-FTLD at Q303, EMD-13708, PDB 7py2^2^. **d. e.** and **f.** TDP-43 fibril in ALS-FTLD at N306, EMD-13708, PDB 7py2^2^. **g. h.** and **i.** TDP-43 fibril in FTLD-TDP type A at Q343, EMD-16628, PDB 8cg3^3^. **j. k.** and **l.** TAF15 fibril in aFTLD at Q60, EMD-16999, PDB 8ons^36^.

**Extended Data Fig. 9.**
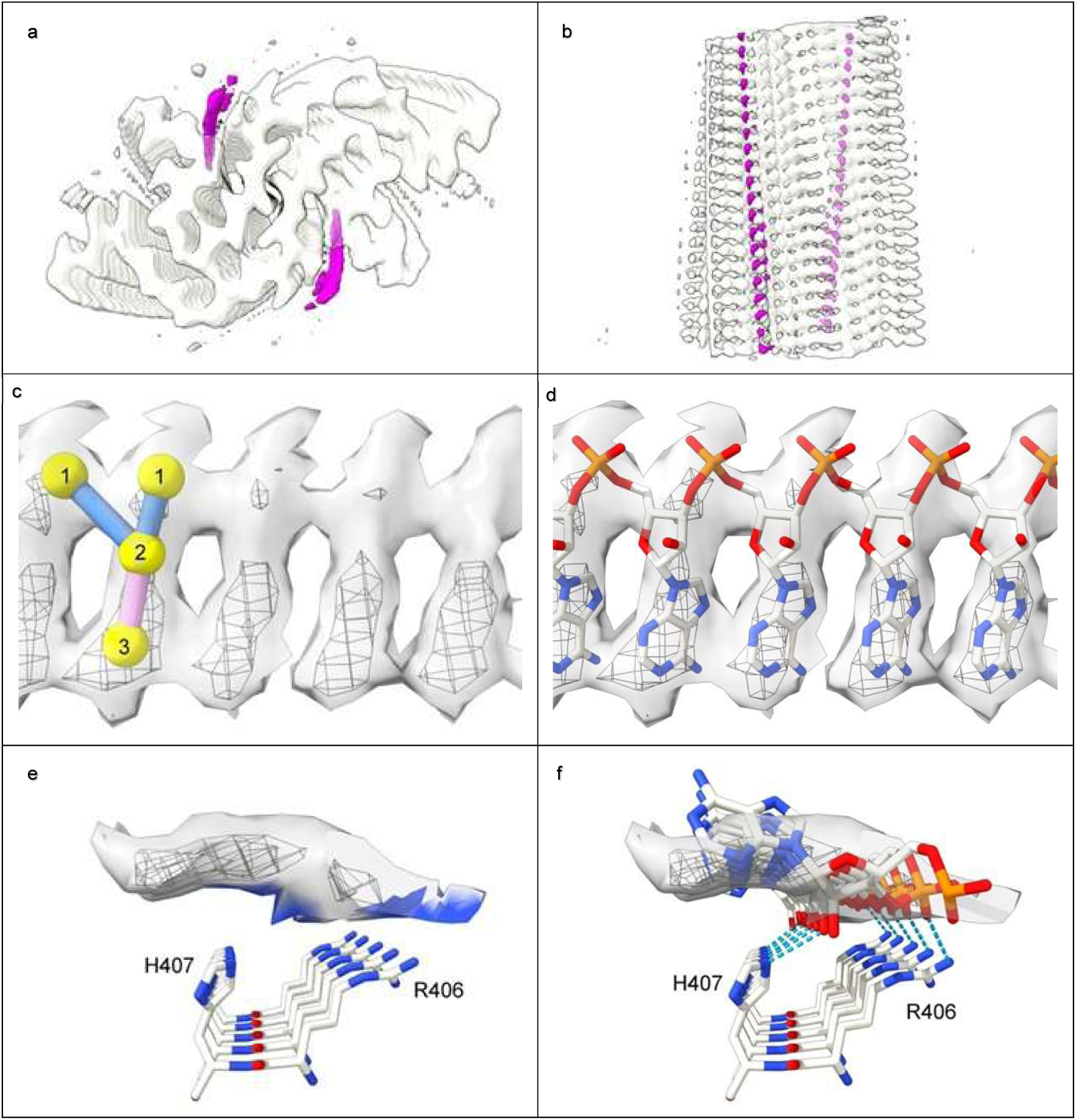
Rod-like densities in RNA-induced tau fibrils ^56^ are similar to LCEDs. **a.** and **b.** Protein white, rod-like extra densities (containing RNA) fuchsia, spacing of protein rungs about 4.8 Å. **c.** and **d.** Extra density at two contour levels, authors’ level in mesh, with markers (**c**) and authors’ model of RNA^56^ (**d**). **e.** and **f.** Extra density in transverse section opposite H407 and R406 basic polar residues, at two levels, authors’ level in mesh. **e.** shows colour zone (blue shading) at 3.5 Å from donor atoms in the protein. **f.** shows authors’ model of RNA^56^ forming hydrogen bonds with donor atoms in histidines and arginines (dashed blue lines). Images of EMD-25364 and PDB 7sp1^56^ created with UCSF ChimeraX^46^.

**Extended Data Table 1.** Cryo-EM maps and LCEDs.

| | All | Aβ | α-syn | PrP | TAF15 | tau | TDP-43 | TMEM106B |
| --- | --- | --- | --- | --- | --- | --- | --- | --- |
| All maps | 161 | 20 | 9 | 2 | 6 | 71 | 9 | 44 |
| Resolution, Å, range (median) | 1.9 to 23 (3.09) | 1.99 to 7.77 (3.07) | 2 to 3.3 (3.09) | 3.13 to 3.29 (3.21) | 1.97 to 2.67 (2.41) | 1.9 to 23 (3.1) | 2.39 to 2.93 (2.68) | 2.39 to 8.8 (3.39) |
| Assessable maps | 141 | 17 | 9 | 2 | 6 | 67 | 8 | 32 |
| Maps with LCEDs of assessable maps (%) | 87/141 (62) | 3/17 (18) | 9/9 (100) | 2/2 (100) | 0/6 (0) | 51/67 (76) | 0/8 (0) | 22/32 (69) |
| LCEDs in assessable maps | 233 | 8 | 15 | 8 | 0 | 158 | 0 | 44 |
| LCEDs per map, range (mean) | 0 to 8 (1.7) | 0 to 4 (0.5) | 1 to 3 (1.7) | 4 to 4 (4) | N/A | 0 to 8 (2.4) | N/A | 0 to 3 (1.4) |
| Strength of LCEDs (% of LCEDs) | s 43.3<br>m 18.9<br>w 37.8 | s 50<br>m 0<br>w 50 | s 33.3<br>m 33.3<br>w 33.3 | s 100<br>m 0<br>w 0 | N/A | s 43<br>m 15.8<br>w 41.1 | N/A | s 36.4<br>m 31.8<br>w 31.8 |
| Morphology of LCEDs (% of LCEDs) | b 12<br>i 4.3<br>t 0.4<br>v 1.3<br>x 30.9<br>y 51.1 | b 25<br>i 0<br>t 0<br>v 0<br>x 25<br>y 50 | b 0<br>i 0<br>t 0<br>v 0<br>x 6.7<br>y 93.3 | b 0<br>i 0<br>t 0<br>v 0<br>x 62.5<br>y 37.5 | N/A | b 13.9<br>i 3.8<br>t 0.6<br>v 0.6<br>x 31<br>y 50 | N/A | b 9.1<br>i 9.1<br>t 0<br>v 4.5<br>x 34.1<br>y 43.2 |
| Protein environment (% of N, where N = LCEDs for which matching atomic models available) | D1 18.0<br>D2 39.3<br>D3 5.2<br>D4 7.6<br>M2 19.4<br>M3 0.5<br>M4 7.1<br>M5 1.4<br>M6 1.4<br>N = 211 | M2 100<br>N = 8 | D1 6.7<br>D2 26.7<br>D3 6.7<br>D4 20<br>M5 20<br>M6 20<br>N = 15 | D1 50<br>M2 50<br>N = 8 | N/A | D1 14.6<br>D2 44.9<br>D3 6.3<br>D4 5.7<br>M2 18.4<br>M3 0.6<br>M4 9.5<br>N = 158 | N/A | D1 45.5<br>D2 36.4<br>D4 18.2<br>N = 22 |

Key: Strength: s = strong, m = moderate, w = weak. Morphology: b = bulky, i = indeterminate, t = three-fold-symmetrical, v = V-shaped, x = X-shaped, y = Y-shaped. Protein Environment: D= pure donor, M= mixed donor and acceptor, 1-6 = number of coordinating residues.
References: All maps<sup>1-44</sup>, Aβ maps<sup>10,13,19,24,35,36,38,40,42</sup>, α-syn maps<sup>27,37,39,41</sup>, PrP maps<sup>17</sup>, TAF15 maps<sup>36</sup>, tau maps<sup>1,5-8,10-12,15,16,19,21-26,29-35,43</sup>, TDP-43 maps<sup>2-4</sup>, TMEM106B maps<sup>5,9,14,18,20,24,28,36,44</sup>

## Notes

### Competing Interest Statement

The authors have declared no competing interest.

## References

1. Arakhamia, T. et al. Posttranslational modifications mediate the structural diversity of tauopathy strains. Cell 180, 633–644.e12 (2020).

2. Arseni, D. et al. Structure of pathological TDP-43 filaments from ALS with FTLD. Nature 601, 139–143 (2022).

3. Arseni, D. et al. TDP-43 forms amyloid filaments with a distinct fold in type A FTLD-TDP. Nature 620, 898–903 (2023).

4. Arseni, D. et al. Heteromeric amyloid filaments of ANXA11 and TDP-43 in FTLD-TDP type C. Nature 634, 662–668 (2024).

5. Chang, A. et al. Homotypic fibrillization of TMEM106B across diverse neurodegenerative diseases. Cell 185, 1346–1355.e15 (2022).

6. Falcon, B. et al. Structures of filaments from Pick’s disease reveal a novel tau protein fold. Nature 561, 137–140 (2018).

7. Falcon, B. et al. Tau filaments from multiple cases of sporadic and inherited Alzheimer’s disease adopt a common fold. Acta Neuropathol 136, 699–708 (2018).

8. Falcon, B. et al. Novel tau filament fold in chronic traumatic encephalopathy encloses hydrophobic molecules. Nature 568, 420–423 (2019).

9. Fan, Y. et al. Generic amyloid fibrillation of TMEM106B in patient with Parkinson’s disease dementia and normal elders. Cell Res 32, 585–588 (2022).

10. Fernandez, A. et al. Cryo-EM structures of amyloid-β and tau filaments in Down syndrome. Nat Struct Mol Biol 31, 903–909 (2024).

11. Fitzpatrick, A. W. P. et al. Cryo-EM structures of tau filaments from Alzheimer’s disease. Nature 547, 185–190 (2017).

12. Fowler, S. L. et al. Tau filaments are tethered within brain extracellular vesicles in Alzheimer’s disease. Nat Neurosci 28, 40–48 (2025).

13. Fu, Z. et al. An electrostatic cluster guides Aβ40 fibril formation in sporadic and Dutch-type cerebral amyloid angiopathy. Journal of Structural Biology 216, 108092 (2024).

14. Ghetti, B. et al. TMEM106B amyloid filaments in the Biondi bodies of ependymal cells. Acta Neuropathol 148, 60 (2024).

15. Ghosh, U. et al. Cryo-EM structures reveal tau filaments from Down syndrome adopt Alzheimer’s disease fold. Acta Neuropathol Commun 12, 94 (2024).

16. Hallinan, G. I. et al. Structure of tau filaments in prion protein amyloidoses. Acta Neuropathol 142, 227–241 (2021).

17. Hallinan, G. I. et al. Cryo-EM structures of prion protein filaments from Gerstmann– Sträussler–Scheinker disease. Acta Neuropathol 144, 509–520 (2022).

18. Hoq, Md. R., et al. Cross-β helical filaments of tau and TMEM106B in gray and white matter of multiple system tauopathy with presenile dementia. Acta Neuropathol 145, 707–710 (2023).

19. Hoq, M. R. et al. Cryo-EM structures of cotton wool plaques’ amyloid β and of tau filaments in dominantly inherited Alzheimer disease. Acta Neuropathol 148, 20 (2024).

20. Jiang, Y. X. et al. Amyloid fibrils in FTLD-TDP are composed of TMEM106B and not TDP-43. Nature 605, 304–309 (2022).

21. Kunach, P. et al. Cryo-EM structure of Alzheimer’s disease tau filaments with PET ligand MK-6240. Nat Commun 15, 8497 (2024).

22. Merz, G. E. et al. Stacked binding of a PET ligand to Alzheimer’s tau paired helical filaments. Nat Commun 14, 3048 (2023).

23. Qi, C. et al. Identical tau filaments in subacute sclerosing panencephalitis and chronic traumatic encephalopathy. Acta Neuropathol Commun 11, 74 (2023).

24. Qi, C. et al. Tau filaments from amyotrophic lateral sclerosis/parkinsonism-dementia complex adopt the CTE fold. Proc. Natl. Acad. Sci. U.S.A. 120, e2306767120 (2023).

25. Qi, C. et al. Tau filaments with the Alzheimer fold in cases with *MAPT* mutations V337M and R406W. bioRxiv 2024.04.29.591661 (2024).

26. Qi, C. et al. Tau filaments with the chronic traumatic encephalopathy fold in a case of vacuolar tauopathy with VCP mutation D395G. Acta Neuropathol 147, 86 (2024).

27. Schweighauser, M. et al. Structures of α-synuclein filaments from multiple system atrophy. Nature 585, 464–469 (2020).

28. Schweighauser, M. et al. Age-dependent formation of TMEM106B amyloid filaments in human brains. Nature 605, 310–314 (2022).

29. Schweighauser, M. et al. Mutation ΔK281 in MAPT causes Pick’s disease. Acta Neuropathol 146, 211–226 (2023).

30. Schweighauser, M. et al. Novel tau filament folds in individuals with *MAPT* mutations P301L and P301T. bioRxiv 2024.08.15.608062 (2024).

31. Seidler, P. M. et al. Structure-based discovery of small molecules that disaggregate Alzheimer’s disease tissue derived tau fibrils in vitro. Nat Commun 13, e5451 (2022).

32. Shi, Y. et al. Cryo-EM structures of tau filaments from Alzheimer’s disease with PET ligand APN-1607. Acta Neuropathol 141, 697–708 (2021).

33. Shi, Y. et al. Structure-based classification of tauopathies. Nature 598, 359–363 (2021).

34. Shi, Y., Ghetti, B., Goedert, M. & Scheres, S. H. W. Cryo-EM structures of chronic traumatic encephalopathy tau filaments with PET ligand flortaucipir. Journal of Molecular Biology 435, 168025 (2023).

35. Stern, A. M. et al. Abundant Aβ fibrils in ultracentrifugal supernatants of aqueous extracts from Alzheimer’s disease brains. Neuron 111, 2012–2020.e4 (2023).

36. Tetter, S. et al. TAF15 amyloid filaments in frontotemporal lobar degeneration. Nature 625, 345–351 (2024).

37. Yan, N. L. et al. Cryo-EM structure of a novel α-synuclein filament subtype from multiple system atrophy. FEBS Letters 599, 33–40 (2025).

38. Yang, Y. et al. Cryo-EM structures of amyloid-β 42 filaments from human brains. Science 375, 167–172 (2022).

39. Yang, Y. et al. Structures of α-synuclein filaments from human brains with Lewy pathology. Nature 610, 791–795 (2022).

40. Yang, Y. et al. Cryo-EM structures of amyloid-β filaments with the Arctic mutation (E22G) from human and mouse brains. Acta Neuropathol (2023).

41. Yang, Y. et al. New SNCA mutation and structures of α-synuclein filaments from juvenile-onset synucleinopathy. Acta Neuropathol 145, 561–572 (2023).

42. Yang, Y. et al. Cryo-EM structures of Aβ40 filaments from the leptomeninges of individuals with Alzheimer’s disease and cerebral amyloid angiopathy. Acta Neuropathol Commun 11, 191 (2023).

43. Zhang, W. et al. Novel tau filament fold in corticobasal degeneration. Nature 580, 283–287 (2020).

44. Zhao, Q. et al. A Tau PET tracer PBB3 binds to TMEM106B amyloid fibril in brain. Cell Discov 10, 50 (2024).

45. Scheres, S. H. W., Ryskeldi-Falcon, B. & Goedert, M. Molecular pathology of neurodegenerative diseases by cryo-EM of amyloids. Nature 621, 701–710 (2023).

46. Pettersen, E. F. et al. UCSF ChimeraX: structure visualization for researchers, educators, and developers. Protein Science 30, 70–82 (2021).

47. Robinson, J. L. et al. Neurodegenerative disease concomitant proteinopathies are prevalent, age-related and APOE4-associated. Brain 141, 2181–2193 (2018).

48. Coulthard, E. J. & Love, S. A broader view of dementia: multiple co-pathologies are the norm. Brain 141, 1894–1897 (2018).

49. Lawson, C. L. et al. EMDataBank unified data resource for 3DEM. Nucleic Acids Res 44, D396–D403 (2016).

50. Berman, H. M. et al. The Protein Data Bank. Acta Crystallogr D Biol Crystallogr 58, 899–907 (2002).

51. Lövestam, S. et al. Assembly of recombinant tau into filaments identical to those of Alzheimer’s disease and chronic traumatic encephalopathy. eLife 11, e76494 (2022).

52. Bridges, L. R. RNA as a component of scrapie fibrils. Sci Rep 14, 5011 (2024).

53. Ginsberg, S. D., Crino, P. B., Lee, V. M.-Y., Eberwine, J. H. & Trojanowski, J. Q. Sequestration of RNA in Alzheimer’s disease neurofibrillary tangles and senile plaques. Ann Neurol. 41, 200–209 (1997).

54. Ginsberg, S. D. et al. RNA sequestration to pathological lesions of neurodegenerative diseases. Acta Neuropathologica 96, 487–494 (1998).

55. Geoghegan, J. C. et al. Selective incorporation of polyanionic molecules into hamster prions. Journal of Biological Chemistry 282, 36341–36353 (2007).

56. Abskharon, R. et al. Cryo-EM structure of RNA-induced tau fibrils reveals a small C-terminal core that may nucleate fibril formation. Proc. Natl. Acad. Sci. U.S.A. 119, e2119952119 (2022).

57. Williams, C. J. et al. MolProbity: more and better reference data for improved all-atom structure validation. Protein Science 27, 293–315 (2018).

58. Watson, J. D. & Crick, F. H. The structure of DNA. Cold Spring Harb Symp Quant Biol 18, 123–131 (1953).

59. Bridges, L. R. RNA as a component of fibrils from Alzheimer’s disease and other neurodegenerations. bioRxiv 2023.02.01.526613 (2023).

60. Oesch, B., Groth, D. F., Prusiner, S. B. & Weissmann, C. Search for a scrapie-specific nucleic acid: a progress report. Ciba Found Symp 135, 209–223 (1988).

61. Narang, H. K. Molecular cloning of single-stranded DNA purified from scrapie-infected hamster brain. Res Virol 144, 375–387 (1993).

62. Telling, G. C. et al. Evidence for the conformation of the pathologic isoform of the prion protein enciphering and propagating prion diversity. Science 274, 2079–2082 (1996).

63. Hardy, J. A. & Higgins, G. A. Alzheimer’s disease: the amyloid cascade hypothesis. Science 256, 184–185 (1992).

64. Pintilie, G. D., Zhang, J., Goddard, T. D., Chiu, W. & Gossard, D. C. Quantitative analysis of cryo-EM density map segmentation by watershed and scale-space filtering, and fitting of structures by alignment to regions. J Struct Biol 170, 427–438 (2010).

65. Schneider, C. A., Rasband, W. S. & Eliceiri, K. W. NIH Image to ImageJ: 25 years of image analysis. Nat Methods 9, 671–675 (2012).

